# Networks that learn the precise timing of event sequences

**DOI:** 10.1101/012195

**Authors:** Alan Veliz-Cuba, Harel Shouval, Krešimir Josić, Zachary P. Kilpatrick

## Abstract

Neuronal circuits can learn and replay firing patterns evoked by sequences of sensory stimuli. After training, a brief cue can trigger a spatiotemporal pattern of neural activity similar to that evoked by a learned stimulus sequence. Network models show that such sequence learning can occur through the shaping of feedforward excitatory connectivity via long term plasticity. Previous models describe how event order can be learned, but they typically do not explain how precise timing can be recalled. We propose a mechanism for learning both the order and precise timing of event sequences. In our recurrent network model, long term plasticity leads to the learning of the sequence, while short term facilitation enables temporally precise replay of events. Learned synaptic weights between populations determine the time necessary for one population to activate another. Long term plasticity adjusts these weights so that the trained event times are matched during playback. While we chose short term facilitation as a time-tracking process, we also demonstrate that other mechanisms, such as spike rate adaptation, can fulfill this role. We also analyze the impact of trial-to-trial variability, showing how observational errors as well as neuronal noise result in variability in learned event times. The dynamics of the playback process determine how stochasticity is inherited in learned sequence timings. Future experiments that characterize such variability can therefore shed light on the neural mechanisms of sequence learning.

## Introduction

Networks of the brain are capable of precisely learning and replaying sequences, accurately representing the timing and order of the constituent events (Conway and Christiansen, 2001; Buhusi and Meck, 2005). Recordings in awake monkeys and rats reveal neural mechanisms that underlie such sequence representation. After a training period consisting of the repeated presentation of a cue followed by a fixed sequence of stimuli, the cue alone can trigger a pattern of neural activity correlated with the activity pattern evoked by the stimulus sequence (Eagleman and Dragoi, 2012; Xu et al., 2012). Importantly, the temporal patterns of the stimulus-driven and cue-evoked activity are closely matched (Shuler and Bear, 2006; Gavornik and Bear, 2014).

Various neural mechanisms have been proposed for learning the duration of a single event (Buonomano, 2000; Rao and Sejnowski, 2001; Durstewitz, 2003; Reutimann et al., 2004; Karmarkar and Buonomano, 2007; Gavornik et al., 2009), as well as the order of events in a sequence (Amari, 1972; Kleinfeld, 1986; Wang and Arbib, 1990; Abbott and Blum, 1996; Jun and Jin, 2007; Fiete et al., 2010; Brea et al., 2013). However, mechanisms for learning the precise timing of multiple events in a sequence remain largely unexplored. The activity of single neurons evolves on the timescale of tens of milliseconds. It is therefore likely that sequences on the timescale of seconds are represented in the activity of populations of cells. Recurrent network architecture could determine activity patterns that arise in the absence of input, but how this architecture can be reshaped by training to support precisely timed sequence replay is not understood.

Long term potentiation (LTP) and long term depression (LTD) are fundamental neural mechanisms that change the strength of connections between neurons (Kandel, 2001). Learning in a wide variety of species, neuron types and parts of the nervous system has been shown to occur through LTP and LTD (Alberini, 2009; Takeuchi et al., 2014; Nabavi et al., 2014). It is therefore natural to ask whether LTP and LTD can play a role in the learning of sequence timing (Karmarkar and Buonomano, 2007; Ivry and Schlerf, 2008), in addition to their proposed role in learning sequence order (Abbott and Blum, 1996; Fiete et al., 2010).

We introduce a neural network model capable of learning the timing of events in a sequence. The connectivity and dynamics in the network are shaped by two mechanisms: long term plasticity and short term facilitation. Long term synaptic plasticity allows the network to encode sequence and timing information in the synaptic weights, while slowly evolving short term facilitation can mark time during event playback. These ideas are quite general, and we show that they do not depend on the particulars of the time-tracking mechanisms we implemented. The impact of stimulus variability and neural noise is largely determined by the trajectory of the time-tracking process. Thus, we predict that errors in event sequence recall may be indicative of the mechanism that encodes them.

## Material and Methods

Our goal was to develop a plastic neural network model that could store arbitrary sequences of event times. To do so we considered interconnected populations whose activity was described in terms of firing rates (Wilson and Cowan, 1972). The synaptic connections between the populations were subject to a rate-based plasticity rule (Dayan and Abbott, 2001): Synaptic weights were increased between coactive neural populations, and weakened if populations were not simultaneously active. Training consisted of sessions during which populations were stimulated one at a time in a specific sequence. Each individual stimulation was of arbitrary duration (Fig. 1B), but the timing in the training pattern was fixed across sessions. After the training period, we cued the first population by providing a short stimulus. If training was successful, this cue triggered a pattern of activity that was close to that evoked by the training sequence. Our aim was to identify a mechanism that allowed the network to learn the order and precise timing of the training sequence (Fig. 1C).

**Figure 1:**
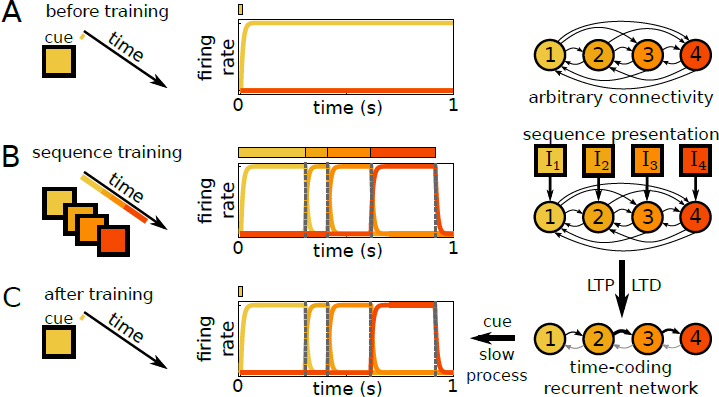
The precise timing of training sequences is learned via long term plasticity. Each stimulus in the sequence is represented by a different color. **A**. Before training, network connectivity is random, and a cue does not trigger a sequential pattern of activity. **B**. During training, a sequence of events is presented repeatedly. Each event activates a corresponding neural population for some amount of time, which is fixed across presentations. Long term plasticity reshapes network architecture to encode the duration and order of these activations. **C**. After sufficient training, a cue triggers the pattern of activity evoked during the training period. Learned synaptic connectivity along with short term facilitation steer activity along the path carved by the training sequence (arrow width and contrast correspond to synaptic strength).

### Population rate model with short term facilitation

Pyramidal cells in cortex form highly connected clusters (Song et al., 2005; Perin et al., 2011), which can correspond to neurons with similar stimulus tuning (Ko et al., 2011). We therefore considered a rate model describing the activity of *N* excitatory populations (clusters), and a single inhibitory population. Here *u*_*j*_ (*j* = 1,…, *N*) represented the activity of different excitatory populations, and *ν* the activity of the inhibitory population. The excitatory and inhibitory populations were coupled via long range connections. We also assumed that population *j* received input *I*_*j*_(*t*), representing an external stimulus. Our model took the form:

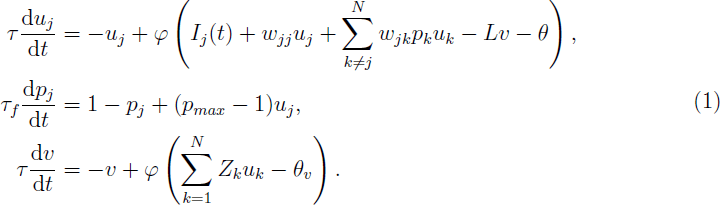

A complete description of the model functions and parameters is given in Table 1. In general, the population firing rates ranged between a small positive value (the background firing rate), and a maximal value (the rate of a driven population), normalized to be 0 and 1, respectively. The baseline weight of the connection from population *k* to *j* was denoted by *w*_*jk*_. Connections within a population were denoted *w*_*jj*_, and these were not subject to short-term facilitation. This assumption did not change our results, but made the analysis more transparent.

Synapses between populations were subject to short term facilitation, and the facilitated connection had weight *w*_*jk*_*p*_*k*_ (Tsodyks et al., 1998). Without loss of generality we considered that short term facilitation varied between 1 and 2 so that the “effective synaptic strength” varied from *w*_*jk*_ to 2*w*_*jk*_. We also assumed *τ*_*f*_ ≫ *τ*, in keeping with the observation that synaptic facilitation dynamics are much slower than changes in firing rates (Markram et al., 1998). Here and below we made a number of other biologically motivated simplifications to make the analysis more transparent. As explained in the Discussion, our results also hold in more detailed models.

In the absence of external stimulus or input from other populations, the dynamics of each population therefore follows 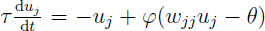. The stationary firing rate is given by the solutions of *φ*(*w*_*jj*_*u*_*j*_ − *θ*) = *u*_*j*_. We typically took *φ* to be a Heaviside step function. Thus, if *w*_*jj*_ > *θ*, there are two equilibrium firing rates: *u*_*j*_ = 0 (inactive population) and *u*_*j*_ = 1 (active population). This assumption simplified the analysis, but was not necessary for our approach to work as shown in the Section **Alternative firing rate response function**.

### Rate-based long term plasticity

We explored how a sequence of inputs presented multiple times to the network could reshape recurrent connectivity so the network replayed the sequence once training was complete. We first describe the unsupervised learning rules that governed the evolution of network connectivity.

Connectivity between the populations in the network was subject to long term potentiation (LTP) and long term depression (LTD). Following experimental evidence (Bliss and Lømo, 1973; Dudek and Bear, 1992; Markram and Tsodyks, 1996; Sjöström et al., 2001), connections were modulated using a rule based on pre- and post-synaptic activity. We also incorporated ‘soft’ bounds to account for the diminishing effects of LTP/LTD as synaptic weights near their maximum/minimum possible values (Gerstner and Kistler, 2002).

We made three main assumptions about the long term evolution of synaptic strength, *w* = *w*_pre→post_: (a) If the presynaptic population activity was low (*u*_*pre*_ ≈ 0), the change in synaptic strength was negligible 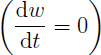; (b) If the presynaptic population was highly active (*u*_*pre*_ = 1) and the postsynaptic population responded weakly (*u*_*post*_ = 0), then the synaptic strength decayed toward zero 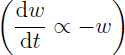; and (c) If both populations had a high level of activity (*u*_*post*_ ≈ 1 and *u*_*pre*_ ≈ 1), then synaptic strength increased toward an upper bound 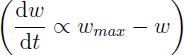. Similar assumptions have been used in previous rate-based models of LTP/LTD (von der Malsburg, 1973; Bienenstock et al., 1982; Oja, 1982; Miller, 1994), and it has been shown that calcium-based (Graupner and Brunel, 2012) and spike-time dependent (Clopath et al., 2010; Gjorgjieva et al., 2011) plasticity rules can be reduced to such rate-based rules (Pfister and Gerstner, 2006). A simple differential equation that implements these assumptions is

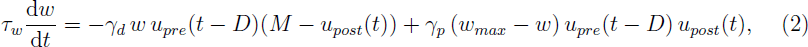

where *τ*_*w*_ is the time scale, *γ*_*d*_ (*γ*_*p*_) represents the strength of LTD (LTP), *D* is a delay accounting for the time it takes for the presynaptic firing rate to trigger plasticity processes, and *M* is a parameter that determines the threshold and magnitude of LTD. Eq. (2) can be derived by linearizing rate-correlation based rules computed from the average of triplet STDP rules (Pfister and Gerstner, 2006). In essence, it is a Hebbian rate-based plasticity rule with soft bounds involving only linear and quadratic dependences of the pre- and post-synaptic rates (Gerstner and Kistler, 2002). Temporal asymmetry that accounts for the causal link between pre- and post-synaptic activity is incorporated with a slight delay in the dependence of pre-synaptic activity *u*_*pre*_(*t* – *D*) (Gütig et al., 2003). This learning rule can also be viewed as a firing-rate version of the calcium-based plasticity model proposed by Graupner and Brunel (2012). Relaxing some of these assumptions leads to qualitatively similar results, as we demonstrate in the Section **Incorporating long timescale plasticity**.

**Table 1:**
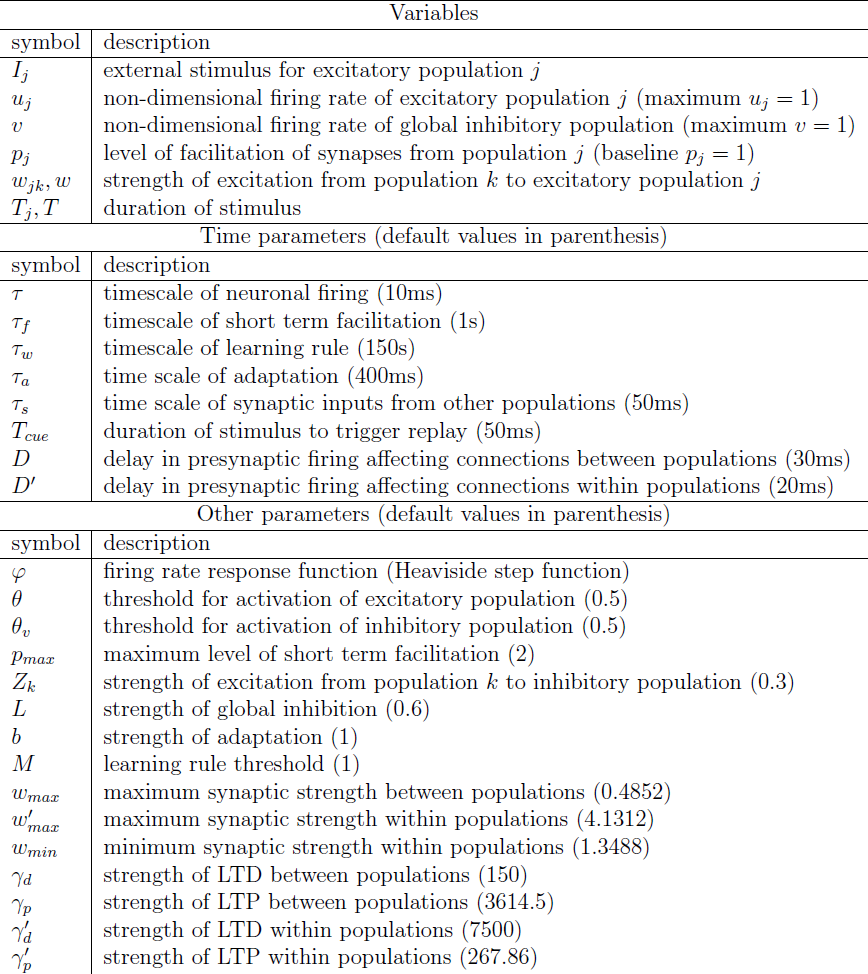
Variables and parameters with their default values. The default values were used in all simulations, unless otherwise noted.

| Variables |  |
| --- | --- |
| symbol | description |
| $I_j$ | external stimulus for excitatory population $j$ |
| $u_j$ | non-dimensional firing rate of excitatory population $j$ (maximum $u_j = 1$) |
| $v$ | non-dimensional firing rate of global inhibitory population (maximum $v = 1$) |
| $p_j$ | level of facilitation of synapses from population $j$ (baseline $p_j = 1$) |
| $w_{jk}, w$ | strength of excitation from population $k$ to excitatory population $j$ |
| $T_j, T$ | duration of stimulus |
| Time parameters (default values in parenthesis) |  |
| symbol | description |
| $\tau$ | timescale of neuronal firing (10ms) |
| $\tau_f$ | timescale of short term facilitation (1s) |
| $\tau_w$ | timescale of learning rule (150s) |
| $\tau_a$ | time scale of adaptation (400ms) |
| $\tau_s$ | time scale of synaptic inputs from other populations (50ms) |
| $T_{cue}$ | duration of stimulus to trigger replay (50ms) |
| $D$ | delay in presynaptic firing affecting connections between populations (30ms) |
| $D'$ | delay in presynaptic firing affecting connections within populations (20ms) |
| Other parameters (default values in parenthesis) |  |
| symbol | description |
| $\varphi$ | firing rate response function (Heaviside step function) |
| $\theta$ | threshold for activation of excitatory population (0.5) |
| $\theta_v$ | threshold for activation of inhibitory population (0.5) |
| $p_{max}$ | maximum level of short term facilitation (2) |
| $Z_k$ | strength of excitation from population $k$ to inhibitory population (0.3) |
| $L$ | strength of global inhibition (0.6) |
| $b$ | strength of adaptation (1) |
| $M$ | learning rule threshold (1) |
| $w_{max}$ | maximum synaptic strength between populations (0.4852) |
| $w'_{max}$ | maximum synaptic strength within populations (4.1312) |
| $w_{min}$ | minimum synaptic strength within populations (1.3488) |
| $\gamma_d$ | strength of LTD between populations (150) |
| $\gamma_p$ | strength of LTP between populations (3614.5) |
| $\gamma'_d$ | strength of LTD within populations (7500) |
| $\gamma'_p$ | strength of LTP within populations (267.86) |

### Encoding timing of event sequences

We next describe the effect of sequential external stimuli (training) on the network model given in Eq. (1) and derive expressions for how the learned synaptic strength evolves depending on the duration of the stimuli.

Our training protocol was based on several recent experiments that explored cortical learning in response to sequences of visual stimuli (Xu et al., 2012; Eagleman and Dragoi, 2012; Gavornik and Bear, 2014). During a training trial, an external stimulus *I*_*j*_(*t*) activated one population at a time. Each individual stimulus could have a different duration (Fig. 1B). We stimulated *n* populations, and enumerated them by order of stimulation; that is, population 1 was stimulated first, then population 2, and so on. This numbering is arbitrary, and the initial recurrent connections have no relation to this order. We denote the duration of input *j* by *T*_*j*_. All inputs stop at *T*_*tot*_ = *T*_1_ + *T*_2_ +…+ *T*_*n*_. A sequence was presented *m* times.

Repeated training of the network described by Eq. (1) with a fixed sequence drove the synaptic weights *w*_*ij*_ to equilibrium values. We assumed that during sequence presentation, the amplitude of external stimuli *I*_*j*_(*t*) was sufficiently strong to dominate the dynamics of the population rates, *u*_*j*_. Then, the activity of the populations during training evolved according to:

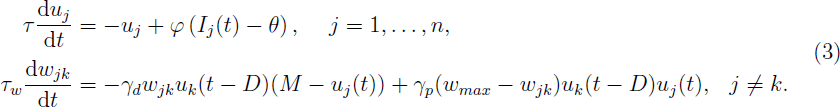

Thus, the timing of population activations mimicked the timing of the input sequence, *i.e.* the training stimulus.

#### Synaptic weights for consecutive activations

For simplicity, we begin by describing the case of two populations, *N* = 2, and we consider the threshold that determines the level of LTD equal to 1, *M* = 1, so that LTD is absent when the postsynaptic population is active (*u*_*post*_ = 1). Suppose that *I*_1_(*t*) = *I*_*S*_ on [0, *T*_1_] and *I*_2_(*t*) = −*I*_*H*_, and *I*_2_(*t*) = *I*_*S*_ on [*T*_1_, *T*_1_ + *T*_2_] and *I*_1_(*t*) = −*I*_*H*_, where *I*_*S*_ and *I*_*H*_ are large enough so that Eq. (3) is valid. We assumed that *T*_*i*_ > *D*, *T*_*i*_ ≫ *τ*, and *τ*_*w*_ ≫ *τ*, so that the stimulus was longer than the plasticity delay, and plasticity slower than the firing rate dynamics. Separation of timescales in Eq. (3) implies that *u*_*j*_ ≈ *φ*(*I*_*j*_(*t*) − *θ*), so the firing rate of populations 1 and 2 is approximated by *u*_1_(*t*) ≈ 1 on [0, *T*_1_] and zero elsewhere, and *u*_2_(*t*) ≈ 1 on [*T*_1_, *T*_1_ + *T*_2_] and zero elsewhere (Fig. 2A). Hence, during a training trial on a time interval [0, *T*_*tot*_], we obtain from Eq. (3) the following piecewise equation for the synaptic strength, *w*_21_, in terms of the duration of the first stimulus, *T*_1_,

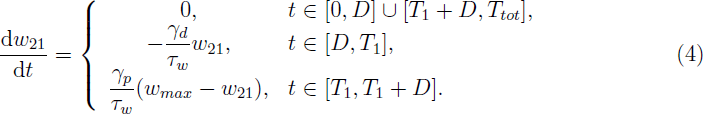

Note that we assumed that *u*_1_(*t*) = 0 for *t* < 0.

Eq. (4) allows the network to encode *T*_1_ using the weight *w*_21_. Namely, solving Eq. (4) we obtain

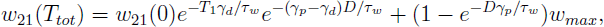

which relates the synaptic weight at the end of a presentation, *w*_21_(*T*_*tot*_), to the synaptic weight at the beginning of the presentation, *w*_21_(0) (See Fig. 2A). Thus, there is a recursive relation that relates the weight *w*_21_ at the end of the *i* + 1st stimulus to the weight at the end of the *i*th stimulus:

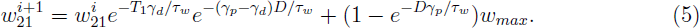

As long as *γ*_*p*_ > *γ*_*d*_, the sequence 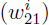 converges to

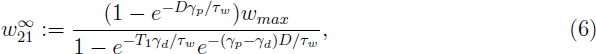

as seen in Fig. 2B. An equivalent expression also holds in the case of an arbitrary number of populations.

The relative distance to the fixed point 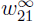 is computed by noting that (for *T*_1_ fixed)

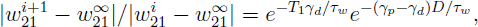

from which we calculate

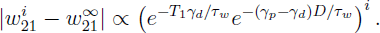

Thus, the sequence converges exponentially with the number of training trials, *i*. The relative distance to the fixed point is proportional to 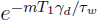, so the convergence is faster for larger values of *T*_1_, Fig. 2C.

**Figure 2:**
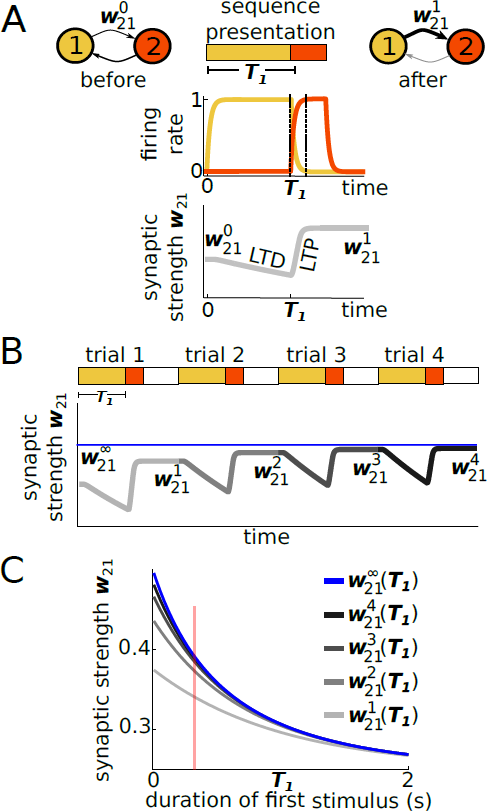
Encoding timing in synaptic weights. **A**. Synaptic connections evolve during training. When a presynaptic population (1) is active and a postsynaptic population (2) is inactive, LTD reduces the synaptic strength *w*_21_. When the populations (1 and 2) are co-active (overlap window between dashed lines), LTP increases *w*_21_. Shortly after, global inhibition inactivates the presynaptic population (1), so long term plasticity ceases (see Materials and Methods). As in the text, 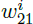 denotes the weight at the end of the *i*-th trial. Arrow width and contrast correspond to synaptic strengths. **B**. After several training trials, the synaptic strength 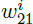 converges to a fixed point, 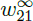, whose amplitude depends on the activation time of the presynaptic population. **C**. Starting from the same initial value, 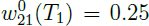, the weight 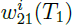 converges to different values, 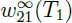, depending on the the training time, *T*_1_. Pink bar at *T*_1_=300ms corresponds to the value used in **A** and **B**.

#### Synaptic weights of populations that are not co-activated

To compute the dynamics of *w*_12_, we note that during a training trial on the time interval [0, *T*_*tot*_], the following piecewise equation governs the change in synaptic weight,

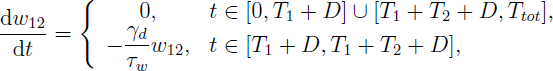

which can be solved explicitly to find

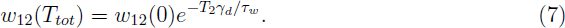

We can therefore write a recursive equation for the weight after the *i* + 1st stimulus in terms of the weight after the *i*th stimulus

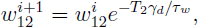

which converges to 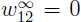. Thus, 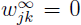 for all pairs of populations (*j*, *k*) for which population *j* was not activated immediately after population *k* during training. In sum, sequential activation of the populations leads to the strengthening only of the weights *w*_*j*+1,*j*_, while other weights are weakened.

### Reactivation of trained networks

We considered a framework based on the experiments of Xu et al. (2012) and Eagleman and Dragoi (2012) who presented a sensory cue at the beginning of each of a number of presentations of a stimulus sequence. After training, the sensory cue was presented to trigger the recall of the training sequence. Gavornik and Bear (2014) employed a related protocol. After repeated presentation of a sequence of sinusoidal gratings, they examined the neural response to the presentation of only part of the sequence. We assumed that synapses were not plastic during replay. As we show in Section **Incorporating long timescale plasticity**, this assumption was not necessary for our approach to work, but it simplified the analytical calculations.

To examine how a sequence of event timings could be encoded by our network, the first neural population in the sequence was activated with a short cue. Typically, this cue was of the form *I*_1_(*t*) = 1 for *t* ∈ [0, *T*_*cue*_], *I*_1_(*t*) = 0 for *t* ∈ [*T*_*cue*_, ∞), and *I*_*j*_(*t*) = 0 for *j* ≠ 1 (Fig. 1C). During replay, aside from the initial cue, the activity in the network was generated through recurrent connectivity.

We describe the case of two populations where the first population is cued, and remains active due to self-excitation (*u*_1_(*t*) ≈ 1), Fig. 3A. Since *u*_2_(0) = 0 and *φ* is the Heaviside step function, the equations governing the dynamics of the second population are

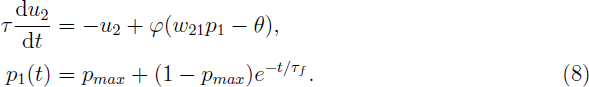

Thus, for population 2 to become active, *w*_21_*p*_1_(*t*) must have reached *θ* (Fig. 3A). The time *T* between when *u*_1_ becomes active and *u*_2_ becomes active (“replay time”) could be controlled by the synaptic strength *w*_21_. Fig. 3B shows the effect of changing the synaptic strength: For very small values of the baseline weight *w*_21_, the facilitating weight *w*_21_*p*_1_(*t*) never reaches *θ* and activation does not occur. Increasing the baseline weight *w*_21_causes more rapid activation of the second population, and for very large weights *w*_21_ the activation is immediate. The weight required for a presynaptic population to activate a postsynaptic population after *T* units of time is given in closed form by

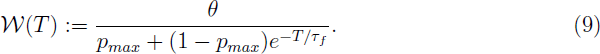

Similarly, the inverse of this function,

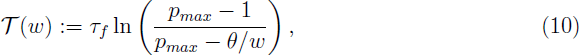

gives the activation time as a function of the synaptic strength (Fig. 3C). Note that Eq. (10) is valid for *θ*/*p*_*max*_ < *w* < *θ*. If *w* ≤ *θ*/*p*_*max*_, then activation of the next population does not occur. If *w* ≥ *θ*, activation is immediate.

To ensure the first population becomes inactive when population 2 becomes active, we assumed that global inhibition overcame the self excitation in the first population, *w*_11_ + *w*_12_*p*_2_(*T*) – *L* − *θ* < 0. Also, for the second population to remain active, we needed the self excitation plus the input received from population 1 to be stronger than the global inhibition; namely, *w*_22_ + *w*_21_*p*_1_(*T*) − *L* − *θ* = *w*_22_ − *L* > 0. These two inequalities are satisfied whenever *w*_12_ is small enough and *L* < *w*_*jj*_ < *L* + *θ*.

### Matching training parameters to reactivation parameters

To guarantee that long term plasticity leads to a proper encoding of event times, it is necessary that the learned weight, 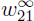 given by Eq. (6), matches the desired weight 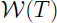 given by Eq. (9). This can be achieved by equating the right hand sides of Eq. (6) and Eq. (9), so that

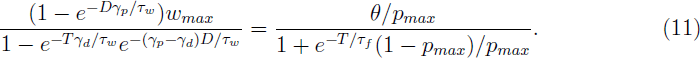

Eq. (11) can be satisfied for all values of *T* by choosing parameters that satisfy

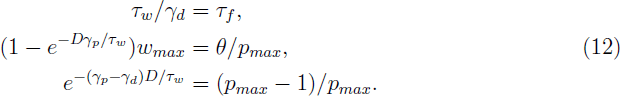

Since there are fewer equations than model parameters, there is a multi-dimensional manifold of parameters for which Eq. (11) holds for all *T*. For instance, for fixed short-term facilitation parameters *θ*, *p*_*max*_, and *τ*_*f*_ and restricting specific plasticity parameters *τ*_*w*_ and *D*, the appropriate *γ*_*d*_, *γ*_*p*_, and *w*_*max*_ can be determined using Eq. (12). This is how we determined the parameters in Figs. 4 and 5.

For more detailed models, the analog of Eq. (12) is more cumbersome or impossible to obtain explicitly. Specifically, when we incorporated noise into our models in the Section **Effect of noise** and considered spike rate adaptation in the Section **An alternative slow process: spike rate adaptation**, we had to use an alternative approach. We found it was always possible to use numerical means to approximate parameter sets that allowed a correspondence between the trained and desired weight for all possible event times. A simple way of finding such parameters was to use the method of least squares: We selected a range of stimulus durations, e.g. [.1*s*, 3*s*], and sampled timings from it, e.g. *S* = {.1*s*, .2*s*, .3*s*,…, 3*s*}. For each *T* ∈ *S* we computed the learned weight, *w*_*learned*_(*T*, pars), where “pars” denotes the list of parameters to be determined. Then, we computed the replay time, *T*_*replay*_(*w*_*learned*_(*T*, pars)). We defined the “cost” function

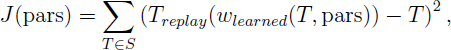

and the desired parameters were given by

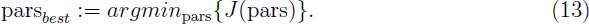

This approach was successful for different models and training protocols, and allowed us to find a working set of parameters for models that included noise or different slow processes for tracking time.

Eq. (13) can also be interpreted as a learning rule for the *network parameters*. Starting with arbitrary network parameter values, any update mechanism that decreases the cost function *J* will result in a network that can accurately replay learned sequence times.

### Training and replay simulations

To test our model, we trained the network with a sequence of four events. Each event in the sequence corresponded to the activation of a single neuronal population (Fig. 4A). Since each population was inactivated (received strong negative input) when the subsequent populations became active, we also assumed that an additional, final population inactivated the population responding to the last event (additional population not shown in the figure). Input during training was strong enough so that activation of the different populations was only determined by the external stimulus overriding global inhibition and recurrent excitation.

During reactivation, the recurrent connections were assumed fixed. This assumption can be relaxed if we assume that LTP/LTD are not immediate, but occur on long timescales, as in the Section **Incorporating long timescale plasticity**.

We used *m* = 10 training trials, the default parameter values in Table 1, and estimated *γ*_*d*_, *γ*_*p*_, and *w*_*max*_ using Eq. (12). After the training trials were finished, we cued the first population in the sequence, using *I*_1_(*t*) = 1 for *t* ∈ [0, *T*_*cue*_] and *I*_*j*_(*t*) = 0 otherwise. We also started with this set of weights, and retrained the network with a novel sequence of stimuli.

### Effect of noise

We also investigated the impact of both stimulus variability and neural noise, focusing on the case of two populations. To examine the impact of variability in stimulus durations, we sampled *T*:= *T*_1_ from a normal distribution (See Fig. 5A) with mean 〈*T*〉 = 0.5 and variance *σ*^2^ = (*c*_*v*_〈*T*〉)^2^, where *c*_*v*_ = 0.1 is the coefficient of variation. We selected 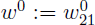 from a uniform distribution on [*θ*/*p*_*max*_, *θ*]. Fig. 5C shows the evolution of the probability density function of *w*^*m*^ as *m* increases (using 20,000 initial *w*^0^’s). We explored the effects of different levels of noise, *σ* = 0.1〈*T*〉, 0.15〈*T*〉, 0.2〈*T*〉 (20,000 initial conditions for each), and estimated the probability density function of *w*^∞^ numerically (Fig. 5E). To see the effect of noise for different mean training durations, we estimated the mean and standard deviation of *w*^∞^ for 〈*T*〉 ranging from 0.1s to 2s (step size of 0.05s) using *σ* = 0.2〈*T*〉 (5,000 initial conditions for each) (Fig. 5G).

We also estimated the mean learned synaptic strength and its variance analytically: Since the synaptic strength, 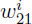, evolved according to the rule

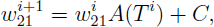

where 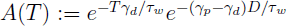 and 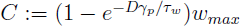, we obtained

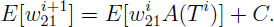

Since 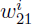 only depends on *T*^*j*^ for *j* < *i*, it follows that 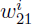 and *A*(*T*^*i*^) are independent random variables; then,

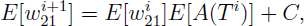

and by taking the limit *i* → ∞ and solving for 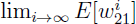 we obtained

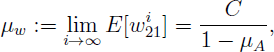

where *μ*_*A*_:= *E*[*A*(*T*^*i*^)]. Squaring 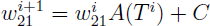 gives

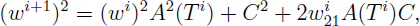

and then it similarly follows that

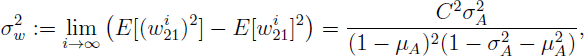

where 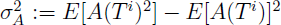. To quantify the average error, we computed numerically the mean and the standard deviation of the replay time (Fig. 5I). The root-mean-square error (RMSE) was computed by

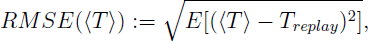

where the expected value is taken over the replay time, *T*_*replay*_.

To introduce neural noise, we added white noise to the rate equations of the populations during training and replay so that

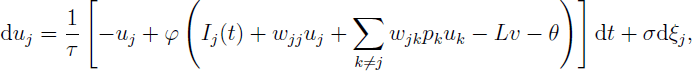

where d*ξ*_*j*_ was a standard white noise process with variance *σ*^2^. The analysis of the effect of noise in population activity is similar to the analysis performed on stimulus duration noise, the only difference being that the noise level was *σ* = 0.03 in panel Fig. 5D, *σ* = 0.03, 0.05, 0.06 in panel Fig. 5F, and *σ* = 0.05 in panels Fig. 5H and 5J.

Note that the parameters given by Eq. (12), which guarantee that training and replay time coincide in the deterministic case, may not be the same as the parameters needed when noise is present. We numerically estimated these parameters using Eq. (13) so the mean training time and mean replay time coincided.

### Alternative firing rate response function

Precise replay relies on a threshold-crossing process which occurs as long as each population is bistable, having only low and high activity states rather than graded activity. Indeed, detailed spiking models (Litwin-Kumar and Doiron, 2012) and experimental recordings (Major and Tank, 2004) suggest that cell assemblies can exhibit multiple stable states. To test whether our conclusions held with different firing rate response functions, we replaced the Heaviside function with the nonsaturating function

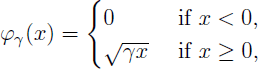

where the parameter *γ* determines the steepness. Note that in the absence of other population inputs,

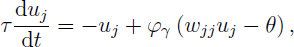

which has steady states determined by the equation *u* = *φ*_*γ*_ (*w*_*jj*_*u* − *θ*). One of the stable steady states is *u* = 0 and there is a positive stable steady state, *u*_*_, which is the largest root of the quadratic equation *u*^2^ − *γw*_*jj*_*u* + *γθ* = 0. For simplicity, we normalized *γ* and self-excitation so that the stable states were *u* = 0 and *u* = 1; namely, we considered 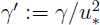 and 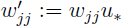.

Since a population is activated when its input reaches the threshold due to short term facilitation, the derivations that led to Eqs. 9 and 10 are still valid for this model. However, the activation of a neuronal population (*u*_*j*_ → 1) was delayed since *φ*_*γ*_ has finite slope. This delay was negligible when firing rate response was modeled by a Heaviside function, and activation was instantaneous. To take this delay into account, we can modify Eqs. 9 and 10 to obtain

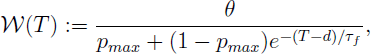

and

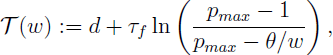

where *d* is a heuristic correction parameter to account for the time it takes for *u*_*j*_ to approach 1.

Following the arguments that led to Eq. (12), we were able to derive constraints on parameters to ensure the correct timings are learned:

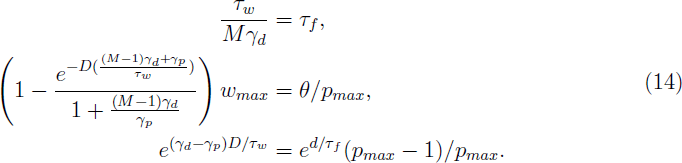

For simulations we used the parameters *M* = 1.5, *d* = 30ms, *γ* = 3, and estimated *γ*_*d*_, *γ*_*p*_, and *w*_*max*_ using Eq. (14) (we then normalized *φ*_*γ*_ and *w*_*jj*_ to make 0 and 1 the stable firing rates). As in the previous simulations, the number of presentations was *m* = 10; the durations of the events were 0.6s, 0.4s, 1s, and 0.5s for events 1, 2, 3 and 4, respectively.

### An alternative slow process: spike rate adaptation

Our results do not depend on the choice of short term facilitation as the time-tracking process. To show this we used spike rate adaptation as an alternative (Benda and Herz, 2003). In contrast to the case of short term facilitation, adaptation causes the effective input from one population to decrease over time.

In this case population activity was modeled by

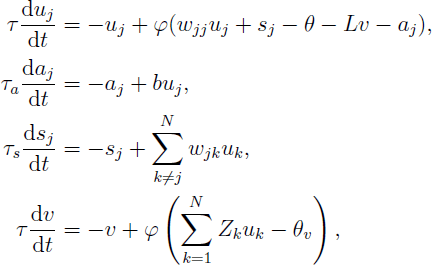

where *a*_*j*_ denotes the adaptation level of population *j*, *τ*_*a*_ is the time scale of adaptation, and *b* is the adaptation strength. Feedback between populations was assumed to be slower than feedback within a population; thus, the total input for population *j* was split into self-excitation (*w*_*jj*_*u*_*j*_), and synaptic inputs from other populations (*s*_*j*_) which evolved on the time scale *τ*_*s*_. Note that in the limit *τ*_*s*_ → 0, synapses are instantaneous.

For a suitable choice of parameters, global inhibition tracks activity faster than excitation between populations. Then, when a population becomes inactive due to adaptation, the level of global inhibition decreases, allowing subsequent populations to become active. This means the strength of self excitation can encode timing. Thus, in this setup we modeled long term plasticity within a population as well. The learning rule for *w*_*jj*_ was analogous to *w*_*jk*_ with the additional assumption that since *w*_*jj*_ represented the synaptic strength within a population, it could not decrease below a certain value *w*_*min*_. Also, the parameters for long term plasticity within a population are allowed to be different from the parameters for long term plasticity between populations.

The learning rule was then

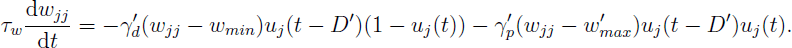

When the population was activated (*u*_1_(*t*) ≈ l) for *t* ∈ [0, *T*_1_], the changes in the weight *w*_11_ were governed by the piecewise differential equation

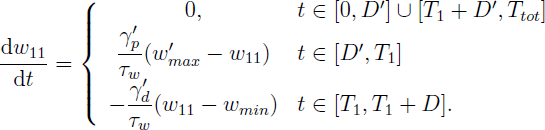

The following equation relates the synaptic weight at the end of a presentation, *w*_11_(*T*_*tot*_), to the synaptic weight at the beginning of the presentation, *w*_11_(0)

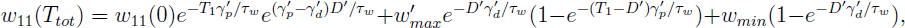

This recurrence relation between the weight at the *i* + 1st stimulus, 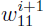, and the weight at the *i*th stimulus, 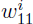, implies that 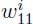 converges to the limit

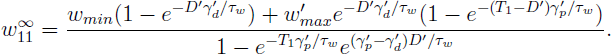

As shown in previous sections, the connection weight between consecutively active populations was strengthened, while weights decreased for other population pairs. Although the connections between populations did not encode time here, they were still used to maintain the order of the sequence. Thus, the precise timing was stored as self excitation strength, and the order of the sequence was encoded as the weights between populations, both of which can be learned.

During replay, if *u*_1_ = 1 we obtain 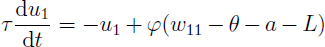, 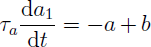. Population 1 will become inactive (*u*_1_ ≈ 0) when *w*_11_ − *a*_1_(*t*) decreases to *θ* + *L*. Then, the next population will become active due to the decrease in global inhibition and the remaining feedback from the first population due to the slower dynamics of feedback between populations.

The precise time of activation can be controlled by tuning the synaptic weight *w*_11_. Furthermore, since the activation time satisfies *w*_11_ − *a*_1_(*T*) = *θ* + *L*, we have a formula that relates the synaptic strength to the activation time.

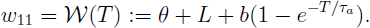

Thus, to guarantee correct time coding and decoding, 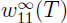 and 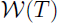 had to be approximately equal for all *T*. The appropriate parameters could not be found in closed form, so we again resorted to finding them numerically using Eq. (13).

For simulations we used the parameters *Z*_*k*_ = 0.6, *L* = 0.8, *γ*_*p*_ = 3750, *γ*_*d*_ = 100, 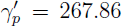, 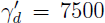, *w*_*max*_ = 1.5, 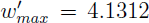, and *w*_*min*_ = 1.3488. As in the previous simulations, the number of presentations was *m* = 10; the duration of the events were 0.6s, 0.4s, 1s, and 0.5s for events 1, 2, 3 and 4, respectively.

### Incorporating long timescale plasticity

Up to this point, we assumed that synaptic changes occurred on the timescale of neural activity. This allowed us to perform a detailed analysis that revealed the main mechanisms underlying time encoding. We also examined a model that incorporated a more detailed description of the synaptic plasticity process. Although this model was less tractable, the main ideas remain unchanged.

Specifically, we addressed the assumption that long term plasticity occurs on the same timescale as the neural activity that initiates it. In addition, we allowed the synaptic connections to change during replay. Previously, we froze synaptic weights during replay, implicitly assuming that a supervisory mechanism could control the prevalence of long term plasticity, depending on whether learning was occurring or not. However, using a more realistic learning rule this assumption can be omitted without changing our results.

Since changes in the synaptic strength are not instantaneous and happen slowly (Alberini, 2009), we considered a model of plasticity separated into two phases: (a) rate correlation detection, which occurs on the timescale of seconds and (b) translation of this information into an actual weight change, which occurs on the timescale of minutes or hours. We refer to the first phase as the updating of *proto-weights* (Gavornik et al., 2009) determined by the pattern of activity in the network. We refer to the second phase as the updating of the *actual* weights, whose evolution depend on the proto-weights. The actual weights exponentially approach the proto-weights, but do so slowly. The model takes the form

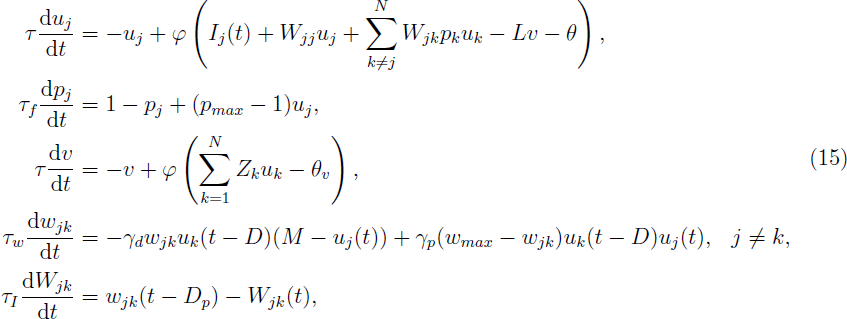

where *τ*_*I*_ is the time scale of the actual weights, and *D*_*p*_ is a time delay in the process of transforming changes in the proto-weights to changes in the actual weights.

It was still possible to analyze the model given in Eqs. 15, but the results were less transparent. During training, the proto-weights satisfied Eq. (5). As the number of training trials increased, the proto-weights converged to the limit given in Eq. (6). The actual weights also slowly approached the same limit; that is, *W*_*jk*_ converged to the limit given in Eq. (6). During replay, the proto-weights evolved according to Eqs. 15, but due to the delay *D*_*p*_ and the time scale *τ*_*I*_, the actual weights remained unchanged and replay occurred accurately as in the Section **Training and replay simulations**.

## Results and statistical analyses

The neural substrates of sequence learning have been explored using a variety of experimental paradigms (Conway and Christiansen, 2001; Ikegaya et al., 2004; Jun and Jin, 2007; Xu et al., 2012; Eagleman and Dragoi, 2012; Gavornik and Bear, 2014) and theoretical approaches (Abbott and Blum, 1996; Wörgötter and Porr, 2005; Karmarkar and Buonomano, 2007; Fiete et al., 2010). Both hippocampal and cortical networks are capable of learning sequences of sensory and motor events (Lee and Wilson, 2002; Janata and Grafton, 2003; Doyon and Benali, 2005). Theoretical models of such networks typically focus on how the correct order of events in a sequence is stored (Abbott and Blum, 1996). However, for many motor (Zatorre et al., 2007) and sensory tasks (Ivry and Schlerf, 2008) learning event timing is as important as learning event order.

Experimental studies of long term memory suggest that learning is associated with changes in the synaptic architecture of neural networks (Bliss et al., 1993; Kandel, 2001; Hofer et al., 2009; Takeuchi et al., 2014; Nabavi et al., 2014). We explored how training-induced synaptic plasticity could allow a neural network to learn the order and precise timing of event sequences. Importantly, we do not assume that a biological clock or rhythm (Treisman, 1963; Miall, 1989; Meck and Benson, 2002) keeps track of elapsed time (see also Karmarkar and Buonomano (2007)).

### Training

We explore sequence learning in a network model of neural populations, where each population is activated by a distinct stimulus or event. The initial connectivity between the populations is random (Fig. 1A). To make our analysis more transparent, we initially consider a deterministic firing rate model (see Materials and Methods). Each individual neural population is bistable, having both a low activity state and a high activity state that is maintained through recurrent excitation. Our results also hold for more biologically plausible firing rate response functions and are robust to noise (see Sections **Other firing rate response functions and slow processes** and **Impact of noise**).

To train the network, we stimulated populations in a fixed order, similar to the training paradigm used by Xu et al. (2012); Eagleman and Dragoi (2012); Gavornik and Bear (2014). The duration of each event in the training sequence was arbitrary (Fig. 1B), and each stimulus in the sequence drove a single neural population. Synaptic connections between populations were plastic. To keep the model tractable, firing rate activity were assumed to immediately alter synaptic connections. Our results extend to a model with synaptic weights changing on longer timescales (see Section **Incorporating non-instantaneous synaptic changes**).

Changes in the network’s synaptic weights depended on the firing rates of the pre- and post-synaptic populations (Bliss and Lømo, 1973; Bienenstock et al., 1982; Dudek and Bear, 1992; Markram and Tsodyks, 1996; Sjöström et al., 2001). When a presynaptic population was active, either: (a) synapses were potentiated (LTP) if the postsynaptic population was subsequently active or (b) synapses were depressed (LTD) if the postsynaptic population was not activated soon after (see Materials and Methods). Such rate-based plasticity rules can be derived from spike time dependent plasticity rules (Kempter et al., 1999; Pfister and Gerstner, 2006; Clopath et al., 2010).

To demonstrate how the timing of events can be encoded in the network architecture, we start with two populations (Fig. 2). During training, population 1 was stimulated for *T*_1_ seconds followed by stimulation of population 2 (Fig. 2A). The stimulus was strong enough to dominate the dynamics of the population responses (see Materials and Methods). While the first stimulus was present, population 1 was active and LTD dominated, decreasing the synaptic weight, *w*_21_, from population 1 to population 2. After *T*_1_ seconds, the first stimulus ended, and the second population was activated. However, population 1 did not become inactive instantaneously, and for some time both population 1 and 2 were active. During this overlap window, LTP dominated leading to an increase in synaptic weight *w*_21_. Shortly after population 1 became inactive, changes in the weight *w*_21_ ceased, as plasticity only occurs when the presynaptic population is active. The initial and final synaptic strengths (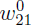 and 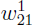, respectively) can be computed in closed form (see Materials and Methods). Repeated presentations of the training sequence leads to exponential convergence of the synaptic strengths, 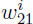 (strength after ith training trial), to a fixed value (Fig. 2B). On the other hand, the synaptic strength *w*_12_ is weakened during each trial because the presynaptic population 2 is always active after the postsynaptic population 1 (see Materials and Methods). In the case of *N* populations, each weight *w*_*k*+1,*k*_ will converge to a nonzero value associated with *T*_*k*_, whereas all other weights will become negligible during replay. Thus, the network’s structure eventually encodes the order of the sequence.

The duration of activation of population 1, *T*_1_, determines the equilibrium value of the synaptic weight from population 1 to population 2, 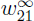 (see Materials and Methods). For larger values of *T*_1_, LTD lasts longer, weakening *w*_21_ (Fig. 2C). Hence, weaker synapses are associated with longer event times. Reciprocally, weaker synapses lead to longer activation times during replay (see Section **A slow process allows precise temporal replay**).

As the stimulus duration, *T*_1_, determines the asymptotic synaptic strength, 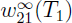, there is a mapping 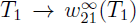 from stimulus times to the resulting weights. Event timing is thus encoded in the asymptotic values of the synaptic strengths.

### A slow process allows precise temporal replay

We next describe how the trained network replays sequences. The presence of a slow process, which we assumed here to be short term facilitation, is critical. This slow process tracks time by *ramping* up until reaching a pre-determined threshold. An event’s duration corresponds to the amount of time it takes the slow variable to reach this threshold. Such ramping models have previously been proposed as mechanisms for time-keeping (Buonomano, 2000; Durstewitz, 2003; Reutimann et al., 2004; Karmarkar and Buonomano, 2007; Gavornik et al., 2009). Without such a slow process, cued activity would result in a sequence replayed in the proper order, but information about event timing would be lost.

For simplicity we focus on two populations, where activity of the first population represents a timed event (Fig. 3). To simplify the analysis, we also assumed that synaptic weights are fixed during replay. This assumption is not essential (see Section **Incorporating non-instantaneous synaptic changes**). After population 1 is activated with a brief cue, it remains active due to recurrent excitation (see Materials and Methods). Meanwhile, short term facilitation leads to an increase in the effective synaptic strength from population 1 to population 2. Population 2 becomes active when the input from population 1 crosses an activation threshold (Fig. 3A). When both populations are simultaneously active, a sufficient amount of global inhibition is recruited to shut off the first population, which receives only weak input from population 2. The second population then remains active, as the strong excitatory input from the first population and recurrent excitation exceed the global inhibition.

**Figure 3:**
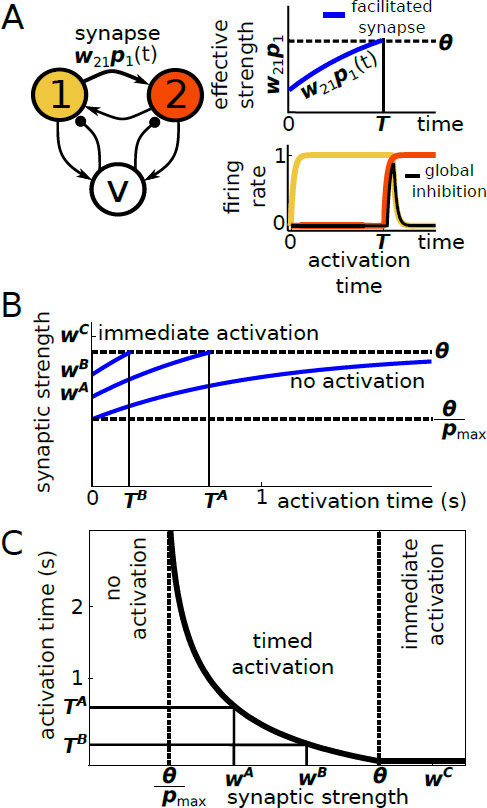
Replay timing. **A**. Once population 1 is actived, the weight of the connection to population 2 slowly increases, eventually becoming strong enough to activate this next population in the sequence. **B**. Activation time *T*:= *T*_1_ decreases monotonically with the weight of the connection between the populations, *w*:= *w*_21_. For weak connections (*w*^*A*^ = 0.33) the synapses must be strongly facilitated to reach the threshold. If the weight is larger (*w*^*B*^ = 0.42) threshold is reached more quickly. For weights above the threshold (*w*^*C*^ = 0.58 > *θ*), population 2 is activated immediately. Activation will not occur when the synaptic connection is smaller than *θ*/*p*_*max*_. **C**. Activation time *T*_1_ plotted against the initial synaptic weight, *w*_21_. For intermediate values of *w*_21_, the relationship is given by Eq. (10). Here *w*^*A*^, *w*^*B*^, *w*^*C*^ and *T*^*A*^, *T*^*B*^ are the same as in panel **B**.

The weight of the connection from population 1 to population 2 determines how long it takes to extinguish the activity in the first population (Fig. 3B). This synaptic weight therefore encodes the time of this first and only event. We demonstrate how this principle extends to multiple event sequences in the Section **Repeated presentation of the same sequence produces a time-coding recurrent network**. The time until the activation of the second population decreases as the initial synaptic strength increases, since a shorter time is needed for facilitation to drive the input from population 1 to the activation threshold (Fig. 3C). Note that when the baseline synaptic strength is too weak, synaptic facilitation saturates before the effective weight reaches the activation threshold, and the subsequent population is never activated. When the baseline synaptic weight is above the activation threshold, the subsequent population is activated instantaneously.

Therefore, long term plasticity allows for encoding an event time in the weight of the connections between populations in the network, while short term facilitation is crucial for replaying the events with the correct timing. The time of activation during cued replay will match the timing in the training sequence as long as training drives the synaptic weights to the value that corresponds to the appropriate event time (Fig. 2C):

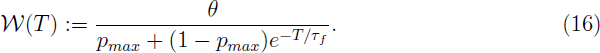

Here *p*_*max*_ and *τ*_*f*_ are the strength and timescale of facilitation (see Materials and Methods). An exact match can be obtained by tuning parameters of the long term plasticity process so the learned weight matches Eq. (16). There is a wide range of parameters for which the match occurs (see Materials and Methods). We next show that the timing and order of sequences containing multiple events can be learned in a similar way.

### Repeated presentation of the same sequence produces a time-coding feedforward network

To demonstrate that the mechanism we discussed extends easily to arbitrary sequences, we consider a concrete sequence of four stimuli. We set the parameters of the model so that the training parameters match the reactivation parameters (see Eqs. 11-13 in Methods).

We trained the network using the event sequence 1-2-3-4 (Fig. 4A), repeatedly stimulating the corresponding populations in succession. The duration of each population activation was fixed across trials. After each training trial, we cued the network by stimulating the first population for a short period of time to trigger replay (see Materials and Methods). Thus, after population 1 is activated the subsequent activity is governed by the network’s architecture. As our theory suggests, the cue-evoked network activity pattern converged with training to the stimulus-driven activity pattern (Fig. 4A).

**Figure 4:**
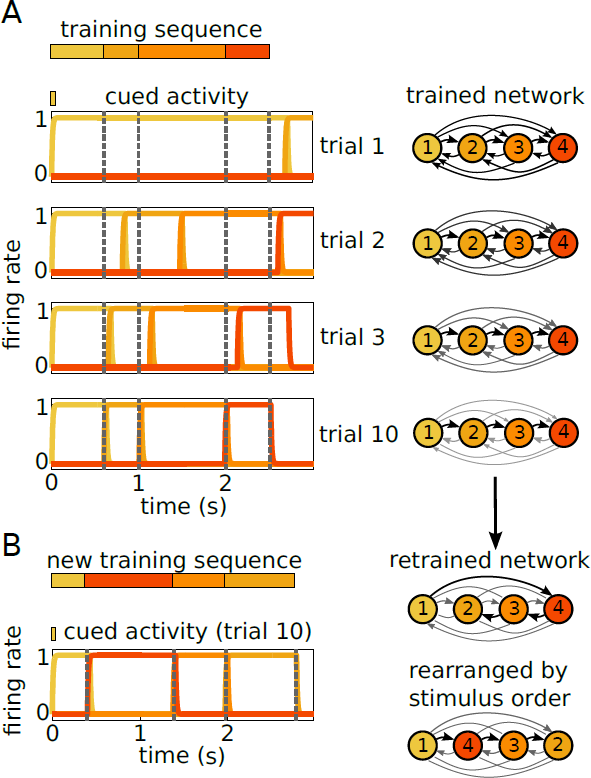
Learning and replaying event sequences. **A**. As the number of training trials increases, a cue results in an activation pattern that approaches that evoked by the training sequence. Network architecture is reshaped to encode the precise duration and order of events in the sequence, with stronger feedforward connections corresponding to shorter events (thicker and darker arrows correspond to larger synaptic strengths). All weights are learned independently and training most strongly affects the weights *w*_*i*→*i*+1_. The event times were *T*_1_ = 0.6, *T*_2_ = 0.4, *T*_3_ = 1, and *T*_4_ = 0.5 seconds, with indices denoting the population stimulated. The initial weights were 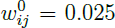 for *i* ≠ *j* and *w*_*ii*_ = 1. **B**. The network in the last row of **A** was retrained with the sequence *T*_1_ = 0.4, *T*_4_ = 1, *T*_3_ = 0.6, and *T*_2_ = 0.8, presented in the order 1-4-3-2. After 10 training trials, the cued network replays the new training sequence.

We further tested whether the same network can be retrained to encode a sequence with a different order of activation (1-4-3-2) with different event times. Fig. 4B shows that after training, the network encodes and replays the new training sequence. Thus, the network architecture can be shaped by long term plasticity to encode an arbitrary sequence of event times, and a brief cue evokes the replay of the learned sequence.

### Impact of noise

To examine the impact of the many sources of variability in the nervous system (Faisal et al., 2008), we explore how noise impacts the training and recall of event sequences in our model. We examine the effects of stochasticity in event times as well as noise in the network activity and the impact such variability has on the training and recall of event sequences. For simplicity, we focus on the case of two populations, but our results extend to sequences of arbitrary length.

To introduce variability in the training times we assumed that the first event time, *T*_1_, was a normally distributed random variable (Fig. 5A), and trained the network as described above. Randomness in the observed event time may be due to variability in the external world, temporal limitations on vision (Butts et al., 2007) or other observational errors (Ma et al., 2006). Event times varied stochastically on each training trial, so learned synaptic strength fluctuated between trials. Thus, we describe the synaptic strength after the *i*th training, 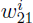, with a probability density function that converges in the limit of many training trials. The peak (mode) of this distribution is the most likely value of the learned synaptic strength after repeated presentation of the sequence (Fig. 5C). The variance of the learned synaptic strength, 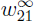, increases monotonically with the variance in the training time, *σ*^2^ (Fig. 5E). In Fig. 5G we show how the probability density function of the learned synaptic strength depends on the mean training time 〈*T*_1_〉. For a distribution of training times with mean 〈*T*_1_〉 we obtain a unimodal probability density for the weights, *p*(*w*_21_). As in the noise-free case, the mode of this weight distribution decreases with 〈*T*_1_〉.

To determine how noise affects activation timing during sequence replay, we compare the mean event time with the mean replayed time. Since the network parameters used here are those obtained from the noise-free case, we expect that replay times are biased. Indeed, Fig. 5I shows that activity during replay is longer on average than the corresponding training event. Also, the variance in activation during replay increases with the mean duration of the trained event. We can search numerically and find a family of parameters for which the mean activation time during replay and training coincide (Fig. 5I, also see Materials and Methods). Error in replay time increases with the duration of the trained time (Fig. 5I, shaded region and inset).

**Figure 5:**
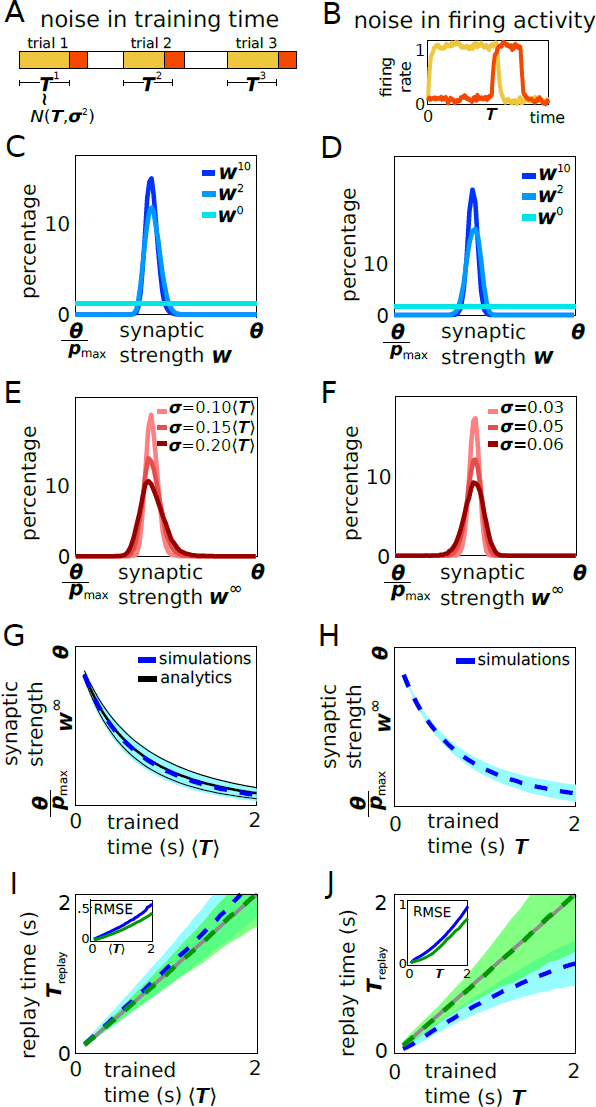
Effect of noise on learning and replay. The effects of adding normally distributed noise to **A** the observed training time *T*:= *T*_1_ (duration of first stimulus) and **B** neural activity *u*_*j*_. Starting from a uniform distribution 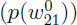 and considering either a noise level of **C** *σ* = 0.1 〈*T*〉 in the training time or **D** *σ* = 0.03 in neural activity, the probability density function 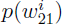 converges to the steady state distribution 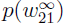 (this is nearly identical to 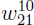). Increasing noise in the training times (**E**) or neural activity (**F**) widens the steady state distribution 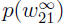. The mean (dashed lines) and standard deviation (shaded region) of 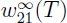 are pictured in panel **G** when noise with standard deviation *σ* = 0.20〈*T*〉 is added to the training times and **H** when noise with standard deviation *σ* = 0.05 is added to neural activity. As in Fig. 2C, the mean learned weight 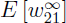 decreases with 〈*T*〉. The mean of the replay time (dashed blue line) and its standard deviation (shaded cyan region) are plotted against 〈*T*〉 for the case of **I** noise in training times and **J** noise in neural populations. As the mean training time 〈*T*〉 increases, so does the effect of noise on replay time, *T*_replay_ (grey lines show the diagonal line *T*_*replay*_ = *T*). For a suitable choice of “corrected-for-noise” parameters, the effect of noise on the mean replay time can be removed (dashed green line and green shaded region). Insets in **I** and **J** show the root-mean-square error in replay time as a function of training time for the parameters obtained from the deterministic case (blue) and corrected-for-noise parameters (green).

We also examined the effect of noise in population firing rates (Fig. 5B). The training and replay protocol were otherwise unchanged. After repeated presentation of a sequence, the distribution of the learned synaptic weights converged (Fig. 5D). The variance of the synaptic strength increased monotonically with the variance of the noise (Fig. 5F), and the mean strength increased monotonically with the event time (Fig. 5H). Since we used the parameters found from the noise-free case, we expect some bias in replay time. After training, the replayed event times are shorter than the corresponding events in the training sequence (Fig. 5J). This systematic bias in the replayed time error is due to the saturating nature of the time-tracking process, short term facilitation (Markram et al., 1998). However, parameters for which mean event time and mean replay time coincide can be found numerically (see Materials and Methods). Also, error in replay time increases with training time duration (Fig. 5I, shaded region and inset).

We found that the network most accurately learns event times that are similar to the timescale of the slow process (Fig. 5H and 5J). Thus, we put forth the testable hypothesis, that networks can learn the precise timing of event sequences by utilizing an inherent slow process whose timescale is similar to the learned event times. To learn event times that are significantly larger or smaller, a different slow processes would be needed.

**Figure 6:**
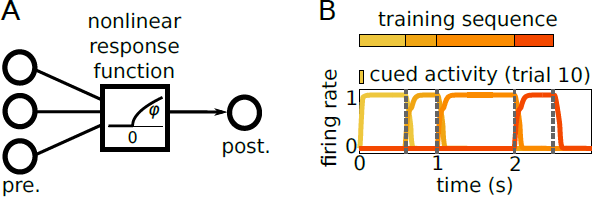
Alternative firing rate function. The mechanism for sequence learning and replay also works for other firing rate functions (see Materials and Methods). **A**. Using a nonlinear and nonsaturating response function 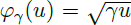, long term plasticity still results in the coding of a training sequence as synaptic weights. **B**. After training, the cued network replays the training sequence similarly to the replay seen in Fig. 4A.

### Other firing rate response functions and slow processes

We next show that the mechanism for learning the precise timing of an event sequence does not depend on the particulars of the model. In previous sections, we used a Heaviside step function as the firing rate function and chose short term facilitation as the slow, time-tracking process. However, the principles we have identified do not depend on these specific choices. More general circuit models of slowly ramping units can learn and replay timed event sequences.

For instance, we can alternatively model the response of a population using a rectified square root 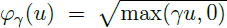, which is nonlinear and nonsaturating (Fourcaud-Trocmé et al., 2003) (Fig. 6A). We can again obtain conditions that guarantee that neural activity during replay matches event duration during training, indicating that our approach works under more realistic assumptions. We use these conditions to find the parameters needed for the network to learn the correct timing, and train the network with the sequence shown in Fig. 4A. Network architecture converges, and the replayed activity matches the order and timing of the training sequence (Fig. 6B).

We also examined whether spike frequency adaptation, *i.e.* a slow decrease in firing rates in response to a fixed input to a neural population, can play the role of a slow, time tracking process (Benda and Herz, 2003), instead of short term facilitation. Long term plasticity can still reshape network architecture so that adaptation allows the network to replay events in the training sequence with the correct timing and the correct order. We explain the mechanism in detail for two populations (Fig. 7 and Materials and Methods).

During training, population 1 is stimulated for *T*_1_ seconds followed by stimulation of population 2 (Fig. 7A). While the first stimulus is present, population 1 is active and LTP dominates, increasing self excitation *w*_11_. After *T*_1_ seconds the stimulus for population 1 ends. When the firing rate of the first population decreases, there is a period of time when LTD dominates and hence *w*_11_ decreases.

Repeated presentations of the training sequence leads to exponential convergence of the synaptic strengths, 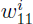, to a fixed value that depends on *T*_1_ (Fig. 7B). Thus, for each stimulus duration a unique synaptic strength is learned. Also, as shown in previous sections, *w*_21_ will converge to a fixed value and *w*_12_ is weakened. Note that in this case timing will be encoded as self excitation strength, and the order will be encoded as the weights between populations.

After training, a cue activates population 1 which then remains active due to recurrent excitation. Meanwhile, adaptation increases and the effective self excitation decreases (Fig. 7C). When adaptation overcomes self excitation, population 1 inactivates. Then, the second population is activated due to the decrease in global inhibition and the remaining feedforward excitation from the first population (see Materials and Methods).

Self excitation within a population determines how long a population remains active, so it serves as an internal memory of time. Furthermore, a wide range of activation times can be encoded using the strength of synaptic self excitation (Fig. 7D). When self excitation is too strong, adaptation will not affect the activity of the first population and deactivation will never occur. On the other hand, when self excitation is too weak, activation is not sustained and the population will be shut off immediately.

This idea generalizes to any number of events and populations. Timing is encoded in the weight of the excitatory self-connections within a population, while sequence order is encoded in the weight of the connections between populations. Moreover, for a range of network parameters, the duration of the sequences during training and reactivation coincide (see Materials and Methods). Presenting the event sequence used in Fig. 4A, the network can learn the precise timing and order of the events (Fig. 7E).

### Incorporating non-instantaneous synaptic changes

We previously assumed that during sequence replay, synaptic connections remained unchanged (see Section **Repeated presentation of the same sequence produces a time-coding recurrent network**). However, if synaptic changes occur on the same timescale as the network’s dynamics, and are allowed to act during replay, the network’s architecture can become unstable. This problem can be solved by assuming that synaptic strengths change slowly compared to network dynamics (Alberini, 2009).

We therefore extended our model so that long term plasticity occurs on more realistic timescales. The impact of rate covariation on the network’s synaptic weights was modeled by a two step process. The initial and immediate signal shaped by the firing rates of pre-and postsynaptic neural populations was modeled by intermediate variables we refer to as *proto-weights* (Gavornik et al., 2009). Changes to the *actual synaptic weights* occur on a much longer timescale, and slowly converged to values determined by the proto-weights (see Materials and Methods).

**Figure 7:**
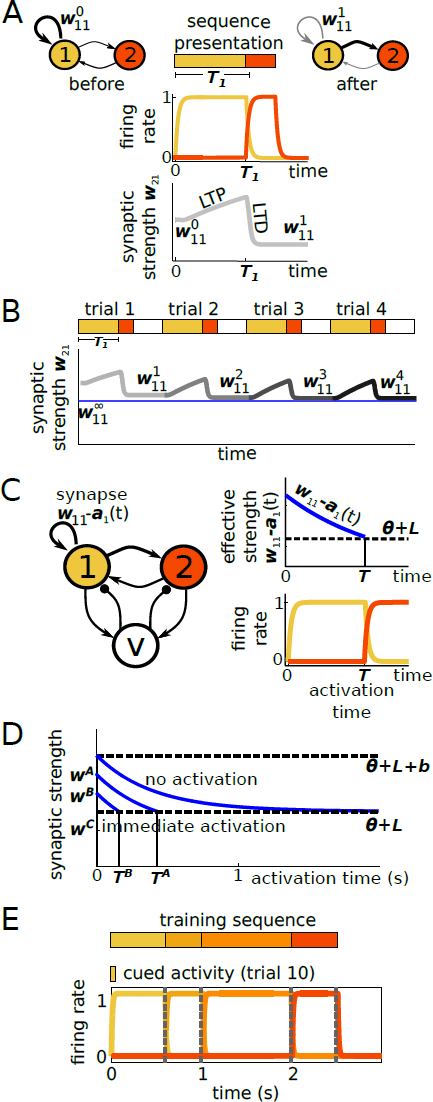
Alternative slow process based on spike frequency adaptation. **A**. When a population is active, LTP increases the synaptic strength *w*_11_. After becoming inactive, LTD decreases *w*_11_ (see Materials and Methods). **B**. Synaptic weight 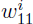 is updated after the *i*th training trial. After several trials, 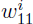 converges to a fixed point 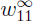 that depends on the activation time of the first population. **C**. Once population 1 is active, adaptation builds up until overcoming self excitation. This will occur when the effective strength, *w*_11_ − *a*_1_(*t*) ([excitation] − [adaptation]), crosses below *θ* + *L* ([threshold] + [inhibition]). **D**. Activation time *T*:= *T*_1_ increases with synaptic strength *w*:= *w*_11_. For strong self excitation (*w*^*A*^) adaptation takes longer to shut off the first population, so the next population in the sequence is activated later. Weaker self excitation (*w*^*B*^) will result in quicker extinction of activity in the first population, result in the next population activating sooner. For self excitation below *θ* + *L* ([threshold] + [inhibition]), the first population will inactivate immediately, resulting in immediate activation of the next population. If self excitation of the first population is greater than *θ* + *L* + *b* ([threshold] + [inhibition] + [maximum adaptation]), it will remain active indefinitely, and the subsequent population is never activated. **E**. When the parameters of the long term plasticity process and the replay process match, the network can learn the precise timing of sequences.

During training, the proto-weights evolve on the same timescale as the neural activity variables (Fig. 8A). Proto-weights evolve on a much faster timescale than the actual weights. Over long times, the actual weights converge to the value of the proto-weights (Fig. 8B), effecting the same synaptic changes as in the previous model. Thus, parameters of the long term plasticity process can still be tuned so that the proper order and timing of event sequences are learned.

Modeling long term plasticity as a two-stage process allowed us to incorporate more realistic details: We were able to assumed that synaptic connections are plastic during replay, as well as during training. Replaying the sequence of neural population activations evokes long term plasticity signals through the proto-weights, and alterations to actual synaptic weights do not take place until after the epochs of neural activity. During reactivation, the actual weights already equal the values necessary to elicit the timed event sequence. Then, the weakening of the connection due to LTD will precisely equal the strengthening due to LTP, resulting in no net change in the actual connection strength. As long as the time scale of synaptic weight consolidation is much larger than the sequence timescale, long term plasticity during replay reinforces the learned network of weights that is already present.

**Figure 8:**
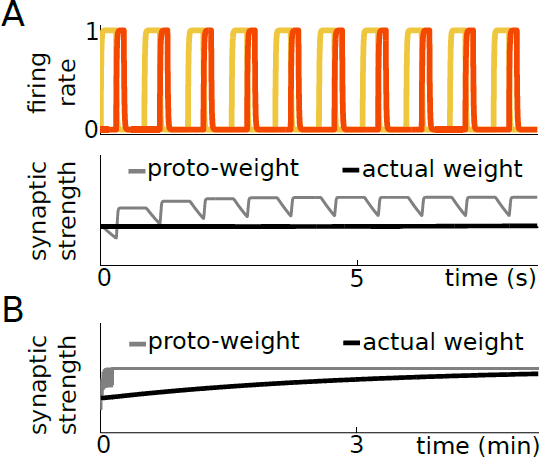
A two-stage learning rule. **A**. *Proto-weights* track the signaled change to synaptic connectivity brought about by firing rate covariance. *Actual weights* evolve on a slower time scale and hence do not track changes in the proto-weights immediately. **B**. Actual synaptic weights approach proto-weight values on a longer timescale, so both eventually converge to the same value.

## Discussion

Sequences of sensory and motor events can be encoded in the architecture of neuronal networks. These sequences can then be replayed with correct order and timing when the first element of the sequence is presented, even in the absence of any other sensory input. Experimental evidence shows that after repeated presentations of a cued training sequence, the presentation of the cue alone triggers a temporal pattern of activity similar to that evoked by the training stimulus (Eagleman and Dragoi, 2012; Xu et al., 2012; Shuler and Bear, 2006; Gavornik and Bear, 2014). Our goal here was to provide a biologically plausible mechanism that could govern the learning of precisely timed sequences.

### Learning both the precise timing and order of sequences

We demonstrated how a complex learning task can be accomplished by combining two simple mechanisms. First, the *timing* of a single event can be represented by a slowly integrating positive feedback process (Buonomano, 2000; Durstewitz, 2003; Reutimann et al., 2004; Shea-Brown et al., 2006; Karmarkar and Buonomano, 2007; Gavornik et al., 2009; Simen et al., 2011). Second, rate dependent long term plasticity can reshape synaptic weights so that the *order* and precise timing of events in a sequence is encoded by the network’s architecture (Amari, 1972; Abbott and Blum, 1996).

To make the problem analytically tractable, we considered an idealized model of neural population firing rates, long term plasticity, and short term facilitation. This allowed us to obtain clear relationships between parameters of the time-tracking process (short term facilitation) and the learning process (long term plasticity). The assumptions about model structure and parameters that were essential for sequence learning could be explicitly described in this model. Similar conditions were required for learning in more realistic models, which incorporated the long timescale of LTP/LTD.

A novel feature of our network model is that long term plasticity influences the length of time a neural population is active. Typical computational models of sequence learning employ networks of neurons (Jun and Jin, 2007; Fiete et al., 2010; Brea et al., 2013) or populations (Abbott and Blum, 1996) that are each active for equal amounts of time during replay. However, sensory and motor processes can be governed by networks whose neurons have a fixed stimulus tuning (Xu et al., 2012; Gavornik and Bear, 2014). Therefore, a sequence of events of varying time lengths should be represented by neural populations that are each active for precisely the length of time of the corresponding event. Our model demonstrates that this can be achieved using rate-based long term plasticity.

### Internal tuning of long term plasticity parameters

There is a large set of parameters for which the network can be trained to accurately replay training sequences. While some parameter tuning is required, in simple cases we could find these parameters explicitly. In all cases appropriate parameters could be obtained computationally using gradient descent. More biologically feasible processes could be employed to find these parameters such as an evolutionary process across many generations, or within an organism whose plasticity processes adapt during development. These are hypothesis that could be experimentally tested. Indeed, recent experimental evidence suggests that networks are capable of internally tuning long term plasticity responses through *metaplasticity* (Abraham, 2008). For instance, NMDA receptor expression can attenuate LTP (Huang et al., 1992; Philpot et al., 2001), while metabotropic glutamate receptor activation can prime a network for future LTP (Oh et al., 2006). We note that such mechanisms would affect the parameters of LTP/LTD, not the synaptic weights themselves.

### Robustness to noise

We have described a mechanism that allows networks to learn sequences in the presence of various sources of noise. Variability in the observation of events is inherited by the encoded sequence timings. As originally proposed in the scalar expectancy theory of Gibbon (1977), errors scale up with the learned timing (Fig. 5I,J). However, experimental observations also reveal that mean recalled interval times closely matched the learned intervals, in both behavioral and neural data (Meck and Malapani, 2004). We found that noise in the observation of the trained times led to mean replayed times that matched the times learned in the absence of any observation error. However, incorporating noise into neural populations led to a shortening of mean replay times, due to the saturation in the ramping process we have chosen. This disparity could be addressed by retuning model parameters to account for this systematic shift in learned timings. An alternative approach could employ a linearly ramping process, which would lead to statistics of replayed times that more closely matched scalar expectancy theory (Durstewitz, 2003; Reutimann et al., 2004; Shea-Brown et al., 2006; Simen et al., 2011).

### Models that utilize ramping processes with different timescales

Our proposed mechanism relies on a ramping process that evolves on the same timescale as the training sequence. Short term facilitation (Markram et al., 1998) as well as rate adaptation (Benda and Herz, 2003) can fulfill this role. However, other ramping processes that occur at the cellular or network level are also capable of marking time. For instance, slow synaptic receptor types such as NMDA can slowly integrate sensory input (Wang, 2002), resulting in population firing rate ramping similar to experimental observations in interval timing tasks (Xu et al., 2014). Were we to incorporate slow recurrent excitatory synapses in this way, the duration of represented events would be determined by the decay timescale of NMDA synapses. Alternatively, we could have also employed short term depression as the slow process in our model. Mutual inhibitory networks with short term depression can represent dominance time durations that depend on the network’s inputs, characteristic of perceptual rivalry statistics (Laing and Chow, 2002). This relationship between population inputs and population activity durations could be leveraged to represent event times in sequences.

Events that occur on much shorter or longer timescales than those we explored here could be marked by processes matched to those timescales. For instance, fast events may be represented simply using synaptic receptors with rapid kinetics, such as AMPA receptors (Clements, 1996). AMPA receptor states evolve on the scale tens of milliseconds, which would allow representation of several fast successive events. Slow events could also be represented by a long chain of sub-populations, each of which is activated for a shorter amount of time than the event. In the context of our model, this would mean each population would contain sub-populations connected as a feedforward chain (Goldman, 2009). Networks of cortical neurons can have different subpopulations with distinct sets of timescales, due to the variety of ion channel and synaptic receptor kinetics (Bernacchia et al., 2011; Pozzorini et al., 2013; Costa et al., 2013). This *reservoir* of timescales could be utilized to learn events whose timings span several orders of magnitude.

### Learning the repeated appearance of an event

We only considered training sequences in which no event appeared more than once (e.g. 1-2-3-4). If events appear multiple times (e.g. 1-2-1-4), then a learned synaptic strength (e.g., *w*_21_) would be weakened when the repeated event appears again. This can be resolved by representing each event repetition by the activation of a different subpopulation of cells. There is evidence that this occurs in hippocampal networks responding to spatial navigation sequences on a figure eight track (Griffin et al., 2007). Even for networks where each stimulus activates a specific population, sequences with repeated stimuli could be encoded in a deeper layer of the underlying sensory or motor system. The same idea can be used to create networks that can store several different event sequences containing the same events (e.g. 1-2-3-4; 2-4-3-1; 4-3-2-1). If multiple sequences begin with the same event (e.g. 1-2-3-4; 1-3-2-4), evoking the correct sequence would require partial stimulation of the sequence (e.g. 1-2 or 1-3). Networks would then be less likely to misinterpret one learned sequence for another sequence with overlapping events (Abbott and Blum, 1996).

### Feedback correction in learned sequences

We emphasize that we did not incorporate any mechanisms for correcting errors in timing during replay. However, this could easily be implemented by considering feedback control via a stimulus that activates the population that is supposed to be active, if any slippage in event timing begins to occur. This assumes there is some external signal indicating how accurately the sequence is being replayed. For instance, human performance of a piece of music relies on auditory feedback signals that are used by the cerebellum to correct motor errors (Zatorre et al., 2007; Kraus and Chandrasekaran, 2010). If feedback is absent or is manipulated, performance deteriorates (Finney and Palmer, 2003; Pfordresher, 2003). Similar principles seem to hold in the replay of visual sequences. Gavornik and Bear (2014) showed that portions of learned sequence are replayed more accurately when preceded by the correct initial portion of the learned sequence. We could incorporate feedback into our model by providing external input to the network at several points in time, not just the initial cue stimulus.

## Conclusions

Overall, our results suggest that a precisely timed sequence of events can be learned by a network with long term synaptic plasticity. Sequence playback can be accomplished by a ramping process whose timescale is similar to the event timescales. Trial-to-trial variability in training and neural activity will be inherited by the sequence representation in a way that depends on the learning process and the playback process. Therefore, errors in sequence representation provide a window into the neural processes that represent them. Future experimental studies of sequence recall that statistically characterize these errors will help to shed light on the neural mechanisms responsible for sequence learning.

## Acknowledgments

We thank Jeffrey Gavornik helpful comments. Funding was provided by NSF-DMS-1311755 (Z.P.K.); NSF/NIGMS-R01GM104974 (A.V-C. and K.J.); and DMS-1122094 (A.V-C. and K.J.).

## Notes

Conflict of interest: We declare no competing interests.

## References

Abbott LF, Blum KI (1996). Functional significance of long-term potentiation for sequence learning and prediction. Cereb Cortex 6:406–416.

Abraham WC (2008). Metaplasticity: tuning synapses and networks for plasticity. Nat Rev Neurosci 9:387.

Alberini C (2009). Transcription factors in long-term memory and synaptic plasticity. Physiol Rev 89:121–145.

Amari SI (1972). Learning patterns and pattern sequences by self-organizing nets of threshold elements. IEEE Trans Comput 21:1197–1206.

Benda J, Herz AV (2003). A universal model for spike-frequency adaptation. Neural Comput 15:2523–2564.

Bernacchia A, Seo H, Lee D, Wang XJ (2011). A reservoir of time constants for memory traces in cortical neurons. Nat Neurosci 14:366–72.

Bienenstock EL, Cooper LN, Munro PW (1982). Theory for the development of neuron selectivity: orientation specificity and binocular interaction in visual cortex. J Neurosci 2:32–48.

Bliss TV, Collingridge GL, et al. (1993). A synaptic model of memory: long-term potentiation in the hippocampus. Nature 361:31–39.

Bliss TVP, Lømo T (1973). Long-lasting potentiation of synaptic transmission in the dentate area of the anaesthetized rabbit following stimulation of the perforant path. J Physiol 232:331–356.

Brea J, Senn W, Pfister JP (2013). Matching recall and storage in sequence learning with spiking neural networks. J Neurosci 33:9565–9575.

Buhusi CV, Meck WH (2005). What makes us tick? functional and neural mechanisms of interval timing. Nat Rev Neurosci 6:755–65.

Buonomano DV (2000). Decoding temporal information: A model based on short-term synaptic plasticity. J Neurosci 20:1129–1141.

Butts DA, Weng C, Jin J, Yeh CI, Lesica NA, Alonso JM, Stanley GB (2007). Temporal precision in the neural code and the timescales of natural vision. Nature 449:92–95.

Clements J (1996). Transmitter timecourse in the synaptic cleft: its role in central synaptic function. Trends Neurosci 19:163–171.

Clopath C, Büsing L, Vasilaki E, Gerstner W (2010). Connectivity reflects coding: a model of voltage-based stdp with homeostasis. Nat Neurosci 13:344–352.

Conway CM, Christiansen MH (2001). Sequential learning in non-human primates. Trends Cogn Sci 5:539–546.

Costa RP, Sjöström PJ, Van Rossum MC (2013). Probabilistic inference of short-term synaptic plasticity in neocortical microcircuits. Frontiers in computational neuroscience 7.

Dayan P, Abbott LF (2001). Theoretical neuroscience. Cambridge, MA: MIT Press.

Doyon J, Benali H (2005). Reorganization and plasticity in the adult brain during learning of motor skills. Curr Opin Neurobiol 15:161–167.

Dudek SM, Bear MF (1992). Homosynaptic long-term depression in area ca1 of hippocampus and effects of n-methyl-d-aspartate receptor blockade. Proc Natl Acad Sci USA 89:4363–4367.

Durstewitz D (2003). Self-organizing neural integrator predicts interval times through climbing activity. J Neurosci 23:5342–5353.

Eagleman SL, Dragoi V (2012). Image sequence reactivation in awake v4 networks. Proc Natl Acad Sci USA 109:19450–19455.

Faisal AA, Selen LP, Wolpert DM (2008). Noise in the nervous system. Nat Rev Neurosci 9:292–303.

Fiete IR, Senn W, Wang CZ, Hahnloser RH (2010). Spike-time-dependent plasticity and heterosynaptic competition organize networks to produce long scale-free sequences of neural activity. Neuron 65:563–576.

Finney S, Palmer C (2003). Auditory feedback and memory for music performance: Sound evidence for an encoding effect. Mem Cognition 31:51–64.

Fourcaud-Trocmé N, Hansel D, Van Vreeswijk C, Brunel N (2003). How spike generation mechanisms determine the neuronal response to fluctuating inputs. J Neurosci 23:11628–11640.

Gavornik JP, Bear MF (2014). Learned spatiotemporal sequence recognition and prediction in primary visual cortex. Nat Neurosci 17:732–737.

Gavornik JP, Shuler MGH, Loewenstein Y, Bear MF, Shouval HZ (2009). Learning reward timing in cortex through reward dependent expression of synaptic plasticity. Proc Natl Acad Sci USA 106:6826–6831.

Gerstner W, Kistler WM (2002). Mathematical formulations of hebbian learning. Biol Cybern 87:404–15.

Gibbon J (1977). Scalar expectancy theory and weber’s law in animal timing. Psychol Rev 84:279.

Gjorgjieva J, Clopath C, Audet J, Pfister JP (2011). A triplet spike-timing–dependent plasticity model generalizes the bienenstock–cooper–munro rule to higher-order spatiotemporal correlations. Proc Natl Acad Sci USA 108:19383–19388.

Goldman MS (2009). Memory without feedback in a neural network. Neuron 61:621–634.

Graupner M, Brunel N (2012). Calcium-based plasticity model explains sensitivity of synaptic changes to spike pattern, rate, and dendritic location. Proc Natl Acad Sci USA 109:3991–3996.

Griffin AL, Eichenbaum H, Hasselmo ME (2007). Spatial representations of hippocampal ca1 neurons are modulated by behavioral context in a hippocampus-dependent memory task. J Neurosci 27:2416–2423.

Gütig R, Aharonov R, Rotter S, Sompolinsky H (2003). Learning input correlations through nonlinear temporally asymmetric hebbian plasticity. The Journal of neuroscience 23:3697–3714.

Hofer SB, Mrsic-Flogel TD, Bonhoeffer T, Hübener M (2009). Experience leaves a lasting structural trace in cortical circuits. Nature 457:313–317.

Huang YY, Colino A, Selig DK, Malenka RC (1992). The influence of prior synaptic activity on the induction of long-term potentiation. Science 255:730–3.

Ikegaya Y, Aaron G, Cossart R, Aronov D, Lampl I, Ferster D, Yuste R (2004). Synfire chains and cortical songs: temporal modules of cortical activity. Science 304:559–564.

Ivry RB, Schlerf JE (2008). Dedicated and intrinsic models of time perception. Trends Cogn Sci 12:273–280.

Janata P, Grafton ST (2003). Swinging in the brain: shared neural substrates for behaviors related to sequencing and music. Nat Neurosci 6:682–687.

Jun JK, Jin DZ (2007). Development of neural circuitry for precise temporal sequences through spontaneous activity, axon remodeling, and synaptic plasticity. PLoS ONE 2:e723.

Kandel ER (2001). The molecular biology of memory storage: a dialogue between genes and synapses. Science 294:1030–1038.

Kappel D, Nessler B, Maass W (2014). Stdp installs in winner-take-all circuits an online approximation to hidden markov model learning. PLoS computational biology 10:e1003511.

Karmarkar UR, Buonomano DV (2007). Timing in the absence of clocks: encoding time in neural network states. Neuron 53:427–438.

Kempter R, Gerstner W, van Hemmen JL (1999). Hebbian learning and spiking neurons. Phys Rev E 59:4498–4514.

Kleinfeld D (1986). Sequential state generation by model neural networks. Proc Natl Acad Sci USA 83:9469–9473.

Ko H, Hofer SB, Pichler B, Buchanan KA, Sjöström PJ, Mrsic-Flogel TD (2011). Functional specificity of local synaptic connections in neocortical networks. Nature 473:87–91.

Kraus N, Chandrasekaran B (2010). Music training for the development of auditory skills. Nat Rev Neurosci 11:599–605.

Laing CR, Chow CC (2002). A spiking neuron model for binocular rivalry. J Comput Neurosci 12:39–53.

Lee AK, Wilson MA (2002). Memory of sequential experience in the hippocampus during slow wave sleep. Neuron 36:1183–1194.

Litwin-Kumar A, Doiron B (2012). Slow dynamics and high variability in balanced cortical networks with clustered connections. Nat Neurosci 15:1498–505.

Ma WJ, Beck JM, Latham PE, Pouget A (2006). Bayesian inference with probabilistic population codes. Nat Neurosci 9:1432–1438.

Major G, Tank D (2004). Persistent neural activity: prevalence and mechanisms. Curr Opin Neurobiol 14:675–84.

Markram H, Tsodyks M (1996). Redistribution of synaptic efficacy between neocortical pyramidal neurons. Nature 382:807–810.

Markram H, Wang Y, Tsodyks M (1998). Differential signaling via the same axon of neocortical pyramidal neurons. Proc Natl Acad Sci USA 95:5323–5328.

Meck WH, Benson AM (2002). Dissecting the brain’s internal clock: how frontal–striatal circuitry keeps time and shifts attention. Brain Cognition 48:195–211.

Meck WH, Malapani C (2004). Neuroimaging of interval timing. Cognitive Brain Res 21:133–137.

Miall C (1989). The storage of time intervals using oscillating neurons. Neural Comput 1:359–371.

Miller KD (1994). A model for the development of simple cell receptive fields and the ordered arrangement of orientation columns through activity-dependent competition between on-and off-center inputs. J Neurosci 14:409–441.

Nabavi S, Fox R, Proulx CD, Lin JY, Tsien RY, Malinow R (2014). Engineering a memory with ltd and ltp. Nature.

Oh MC, Derkach VA, Guire ES, Soderling TR (2006). Extrasynaptic membrane trafficking regulated by glur1 serine 845 phosphorylation primes ampa receptors for long-term potentiation. J Biol Chem 281:752–8.

Oja E (1982). Simplified neuron model as a principal component analyzer. J Math Biol 15:267–273.

Perin R, Berger TK, Markram H (2011). A synaptic organizing principle for cortical neuronal groups. Proc Natl Acad Sci USA 108:5419–5424.

Pfister JP, Gerstner W (2006). Triplets of spikes in a model of spike timing-dependent plasticity. J Neurosci 26:9673–9682.

Pfordresher PQ (2003). Auditory feedback in music performance: Evidence for a dissociation of sequencing and timing. Journal of Experimental Psychology: Human Perception and Performance 29:949.

Philpot BD, Sekhar AK, Shouval HZ, Bear MF (2001). Visual experience and deprivation bidirectionally modify the composition and function of NMDA receptors in visual cortex. Neuron 29:157–169.

Pozzorini C, Naud R, Mensi S, Gerstner W (2013). Temporal whitening by power-law adaptation in neocortical neurons. Nature neuroscience 16:942–948.

Rao RP, Sejnowski TJ (2001). Spike-timing-dependent hebbian plasticity as temporal difference learning. Neural Comput 13:2221–2237.

Reutimann J, Yakovlev V, Fusi S, Senn W (2004). Climbing neuronal activity as an event-based cortical representation of time. J Neurosci 24:3295–3303.

Shea-Brown E, Rinzel J, Rakitin BC, Malapani C (2006). A firing rate model of parkinsonian deficits in interval timing. Brain Res 1070:189–201.

Shuler MG, Bear MF (2006). Reward timing in the primary visual cortex. Science 311:1606–1609.

Simen P, Balci F, Cohen JD, Holmes P, et al. (2011). A model of interval timing by neural integration. J Neurosci 31:9238–9253.

Sjöström PJ, Turrigiano GG, Nelson SB (2001). Rate, timing, and cooperativity jointly determine cortical synaptic plasticity. Neuron 32:1149–1164.

Song S, Sjöström PJ, Reigl M, Nelson S, Chklovskii DB (2005). Highly nonrandom features of synaptic connectivity in local cortical circuits. PLoS Biol 3:e68.

Takeuchi T, Duszkiewicz AJ, Morris RG (2014). The synaptic plasticity and memory hypothesis: encoding, storage and persistence. Philos T Roy Soc Lond B 369:20130288.

Treisman M (1963). Temporal discrimination and the indifference interval: Implications for a model of the “internal clock”. Psychol Monogr-Gen A 77:1.

Tsodyks M, Pawelzik K, Markram H (1998). Neural networks with dynamic synapses. Neural Comput 10:821–835.

von der Malsburg C (1973). Self-organization of orientation sensitive cells in the striate cortex. Kybernetik 14:85–100.

Wang D, Arbib M (1990). Complex temporal sequence learning based on short-term memory. P IEEE 78:1536–1543.

Wang XJ (2002). Probabilistic decision making by slow reverberation in cortical circuits. Neuron 36:955–68.

Wilson HR, Cowan JD (1972). Excitatory and inhibitory interactions in localized populations of model neurons. Biophys J 12:1–24.

Wörgötter F, Porr B (2005). Temporal sequence learning, prediction, and control: a review of different models and their relation to biological mechanisms. Neural Comput 17:245–319.

Xu M, Zhang SY, Dan Y, Poo Mm (2014). Representation of interval timing by temporally scalable firing patterns in rat prefrontal cortex. Proc Natl Acad Sci USA 111:480–485.

Xu S, Jiang W, Poo Mm, Dan Y (2012). Activity recall in a visual cortical ensemble. Nat Neurosci 15:449–455.

Zatorre RJ, Chen JL, Penhune VB (2007). When the brain plays music: auditory–motor interactions in music perception and production. Nat Rev Neurosci 8:547–558.

